# Mapping protein–exopolysaccharide binding interaction in *Staphylococcus epidermidis* biofilms by live cell proximity labeling

**DOI:** 10.1101/2023.08.29.555326

**Authors:** Luan H. Vo, Steven Hong, Kaitlyn E. Stepler, Sureshee M. Liyanaarachchi, Jack Yang, Peter Nemes, Myles B. Poulin

## Abstract

Bacterial biofilms consist of cells encased in an extracellular polymeric substance (EPS) composed of exopolysaccharides, extracellular DNA, and proteins that are critical for cell–cell adhesion and protect the cells from environmental stress, antibiotic treatments, and the host immune response. Degrading EPS components or blocking their production have emerged as promising strategies for prevention or dispersal of bacterial biofilms, but we still have little information about the specific biomolecular interactions that occur between cells and EPS components and how those interactions contribute to biofilm production. *Staphylococcus epidermidis* is a leading cause of nosocomial infections as a result of producing biofilms that use the exopolysaccharide poly- (1→6)-β-*N*-acetylglucosamine (PNAG) as a major structural component. In this study, we have developed a live cell proximity labeling approach combined with quantitative mass spectrometry-based proteomics to map the PNAG interactome of live *S. epidermidis* biofilms. Through these measurements we discovered elastin-binding protein (EbpS) as a major PNAG-interacting protein. Using live cell binding measurements, we found that the lysin motif (LysM) domain of EbpS specifically binds to PNAG present in *S. epidermidis* biofilms. Our work provides a novel method for the rapid identification of exopolysaccharide-binding proteins in live biofilms that will help to extend our understanding of the biomolecular interactions that are required for bacterial biofilm formation.

## Introduction

Biofilms are multicellular communities of microbial cells embedded within a self-produced extracellular polymeric substance (EPS) that is composed largely of extracellular DNA, cell surface and secreted proteins, and exopolysaccharides.^1,2^ The EPS serves as a protective barrier, shielding the bacteria from environmental stressors,^3^ host immune defenses,^4,5^ and antibiotic treatment,^6^ while also facilitating cell–cell and cell–surface adhesion.^2,7–9^ As a result of this adhesion, biofilms are a frequent cause of persistent and nosocomial infections,^3^ in particular those associated with implanted medical devices. Within the EPS, exopolysaccharides and proteins are thought to play the primary role in cell–cell and cell–surface adhesion, but very little information is available about how these molecules interact to facilitate adhesion.^9,10^

Several structurally distinct exopolysaccharides have been associated with biofilm formation of different bacterial species,^10^ and include highly anionic polysaccharides like alginate,^11^ neutral polysaccharides like cellulose,^12^ Psl,^13^ and vibrio polysaccharide (VPS),^14^ and cationic polysaccharides including partially de-*N*-acetylated Pel^15,16^ and poly-(1→6)-β-*N*-acetylglucosamine (PNAG).^17–19^ Of these, PNAG is of particular interest due to its prevalence in the biofilms of common human pathogens, including Gram-positive bacteria *Staphylococcus epidermidis* and *Staphylococcus aureus*,^17,20,21^ and Gram-negative bacteria like *Escherichia coli*,^19^ *Klebsiella pneumoniae*,^22^ and *Acinetobacter baumanii*.^23^ Despite the widespread distribution of PNAG in microbial pathogens and its role in biofilm assembly,^24^ there is still little known about the biomolecular interactions of PNAG that facilitate cell adhesion and biofilm formation.

It is widely thought that the cationic charge of PNAG facilitates cell–cell adhesion by masking the anionic charge of bacterial cell surface components like lipopolysaccharide (LPS) or wall teichoic acids (WTA).^4,25^ However, work from the Pier lab reports that WTA, the most abundant polyanion of the Gram-positive cell envelope, are not required for anchoring PNAG to the cell surface of *S. aureus* cells,^26^ suggesting other biomolecular interactions are required. Recently, there have been several reports of exopolysaccharide–protein interactions that contribute to cell–cell adhesion during biofilm formation.^7,27–31^ For example, binding between LecB and CdrA to the Psl polysaccharide in *P. aeruginosa* biofilms contribute to cell–cell junction formation,^7,27,28^ and binding of RbmA, RbmC, and Bap1 to VPS contribute to biofilm formation in *Vibrio cholerae*.^29–31^ However, reports of specific protein–exopolysaccharide binding interactions within bacterial biofilms are rare, and in the case of PNAG specifically, no protein– PNAG binding interactions have been reported to date.

The lack of experimental evidence for exopolysaccharide-binding proteins in bacterial biofilms is due, at least in part, to a lack of general tools available to capture unknown protein– carbohydrate binding interactions within live biofilm samples. Capturing protein–carbohydrate binding interactions is particularly challenging as binding interactions between individual proteins and carbohydrates are often transient and weak (i.e., mM binding affinity), which rely on multivalent display of the protein and/or carbohydrate structure on the cell surface to increase avidity.^32^ This difficulty is additionally compounded within bacterial biofilms as the methods required to disrupt the biofilm EPS to isolate the proteins require harsh treatment that disrupt weak protein–exopolysaccharide binding interactions. Thus, there is a need for general approaches that would enable the identifications of exopolysaccharide-binding proteins in live bacterial biofilms. Here we report the development of a live cell proximity-labeling approach that enables biotinylation of proteins based on proximity to PNAG within live bacterial biofilms. The approach uses a catalytically inactive PNAG glycosyl hydrolase enzyme Dispersin B (DspB) mutant fused to the engineered ascorbate peroxidase APEX2^33^ to bind PNAG in live bacterial biofilms and catalyze the proximity labeling of PNAG-interacting proteins. Using biofilms of *S. epidermidis* RP62A, we demonstrate that this proximity labeling is PNAG dependent. Combining the PNAG-dependent proximity labeling of live biofilms with quantitative high-resolution mass spectrometry (HRMS)-based proteomics profiling enabled the identification of EbpS as an endogenous *S. epidermidis* PNAG-binding protein.

## Results and discussion

### Proximity labeling of PNAG interactors in *S. epidermidis* biofilms

To generate a fusion protein suitable for tagging proteins in proximity to PNAG within live biofilms, we fused the engineered ascorbate peroxidase enzyme APEX2^33,34^ to a catalytically inactive DspB_E184Q_ mutant through a flexible GSGGGGS linker (Fig. 1A), hereafter referred to as APEX2-DspB. This DspB_E184Q_ mutant has been shown to lack PNAG hydrolase activity^35–37^ but retains the ability to bind PNAG and has previously been genetically fused to enhanced green fluorescent protein (eGFP) for fluorescent labeling of PNAG within live biofilms.^38,39^ APEX2 was chosen over other peroxidase enzymes due to its small size and monomeric nature, improving the likelihood that the fusion protein would effectively penetrate and bind to PNAG within *S. epidermidis* biofilms. The fusion protein was efficiently prepared and purified from recombinant *E. coli* (SI Fig.1A). Heme loading of recombinant APEX2-DspB was estimated using the ratio of the Soret band of heme at 405 nm to the protein absorbance at 280 nm, and protein having a ratio < 1 was reconstituted with heme *in vitro* to ensure high heme loading (SI Fig. 1B).

**Figure 1.**
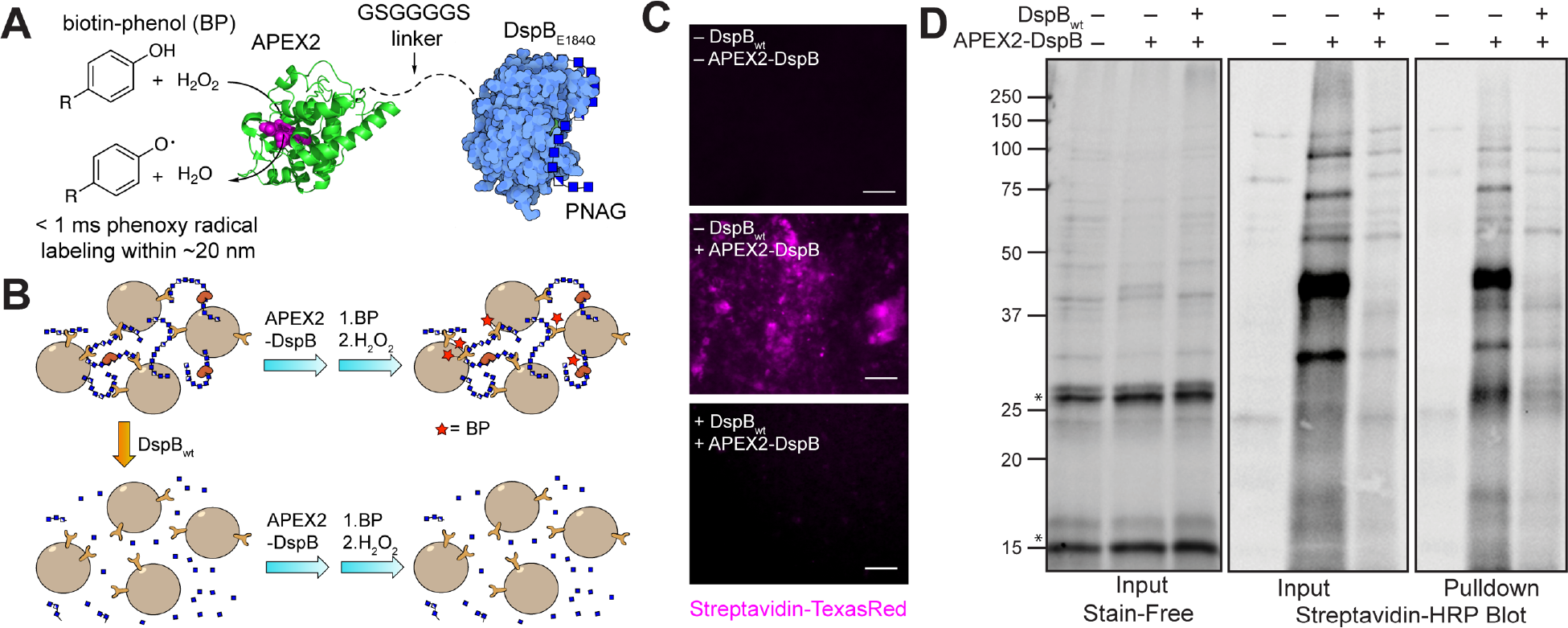
Design and evaluation of APEX2-DspB fusion proteins for live cell proximity labeling of the PNAG interactome in live *S. epidermidis* RP62A biofilms. (A) Schematic of the fusion construct containing an *N*-terminal APEX2 domain connected via a flexible GSGGGGS linker to a catalytically inactive *A. actinomycetemcomitans* DspB_E184Q_ mutant. APEX2 is a heme-dependent ascorbate peroxidase enzyme that has been engineered to catalyze the oxidation of phenol substrates (ie. biotin-phenol, BP) using H_2_O_2_ to generate short lived (< 1 ms) phenoxy radicals. (B) Outline of a live cell proximity labeling experiment. Replicate *S. epidermidis* 48 hr. biofilm samples are incubated with either either APEX2-DspB alone (– +) or in the presence of DspB_wt_ (+ +) for 30 min to digest any PNAG in the sample. Control cells incubated with buffer alone (– –) were also measured in parallel. All samples were incubated with BP (2.5 mM) for 30 min followed by H_2_O_2_ (0.5 mM) for 1 min for sample labeling. (C) Fluorescent microscopy imaging of biotin-labeled *S. epidermidis* biofilm samples imaged using a streptavidin-TexasRed conjugate showing significantly more biotin-labeling in the presence of APEX2-DspB alone (– +) than in control samples incubated with buffer alone (– –), or samples treated with both APEX2-DspB and DspB_wt_ to degrade PNAG (+ +) (scale bar = 20 µm). (D) A representative pull-down experiment and Western blot used to detect biotinylated proteins present after live cell proximity labeling using streptavidin-HRP to visualize biotin-labeled proteins. Total protein content (right), biotin-labeled protein (center) and proteins eluted from pull-down experiments with streptavidin-coated magnetic beads (left) are shown. Asterisks (*) indicate the presence of lysostaphin (∼27 kDa) and hen egg white lysozyme (∼15 kDa) included in the cell disruption buffer.

To evaluate whether APEX2-DspB would be suitable for labeling PNAG-interacting proteins when added exogenously to live *S. epidermidis* biofilms, we incubated 48 h old static *S. epidermidis* RP62A biofilm grown in 6-well microtiter plates with the recombinant protein (Fig. 1B). Biofilm samples were incubated with either buffer alone (– –), 2 µM APEX2-DspB (– +), or a combination of 2 µM wild-type dispersin B (DspB_wt_) and 2 µM APEX2-DspB (+ +). DspB_wt_ was used to enzymatically hydrolyze PNAG within the *S. epidermidis* biofilms prior to labeling and was used to determine if the proximity labeling observed is dependent on PNAG. After washing the cells in buffer, stepwise addition of biotin-phenol and H_2_O_2_, and quenching with ascorbate and Trolox, the biotinylated biofilms were visualized by fluorescent microscopy using a streptavidin-TexasRed probe (Fig. 1C). We observed significant fluorescent labeling in (– +) samples, but no significant fluorescence in either control (– –) samples or (+ +) samples in which PNAG was hydrolyzed by DspB_wt_

To verify that these signals are the result of proximity biotinylation of proteins present with the *S. epidermidis* biofilms, the total protein was isolated from the biofilm samples after proximity labeling. We pulled down biotinylated proteins with streptavidin-coated magnetic beads and then analyzed by Western blot as shown in Fig. 1D. The biotin-labeled proteins were visualized using a streptavidin-horseradish peroxidase (HRP) conjugate. We observed substantially more protein labeling in both the input and pull-down fractions from (– +) samples than from (– –) or (+ +) biofilm samples.

### Identification of Biotinylated PNAG-Interacting Proteins using Quantitative MS-Based Proteomics

To identify the biotinylated proteins observed by Western blot, 48-h-old *S. epidermidis* biofilms were subjected to the same proximity labeling procedure described above, followed by cell lysis, streptavidin pull-down, and on-bead trypsin digestion (Fig. 2A). Three biological replicates of *S. epidermidis* biofilms were subjected to this procedure, and the resulting peptides from each sample were individually labeled with tandem mass tags (TMTs), thus chemically labeling free amines and the *N*-termini of the peptides with a unique isobaric tag that can be identified with tandem MS (MS^2^) and quantified at the MS^3^ stage.^40–42^ The TMT-tagged peptides from all nine samples (3 biological replicates labeled using the 3 different proximity labeling conditions) were combined and analyzed as a single multiplexed sample by nano-flow high-performance liquid chromatography coupled with high-resolution multistage mass spectrometry (nanoLC-MS^n^), ensuring accurate comparison of protein concentrations.

**Figure 2.**
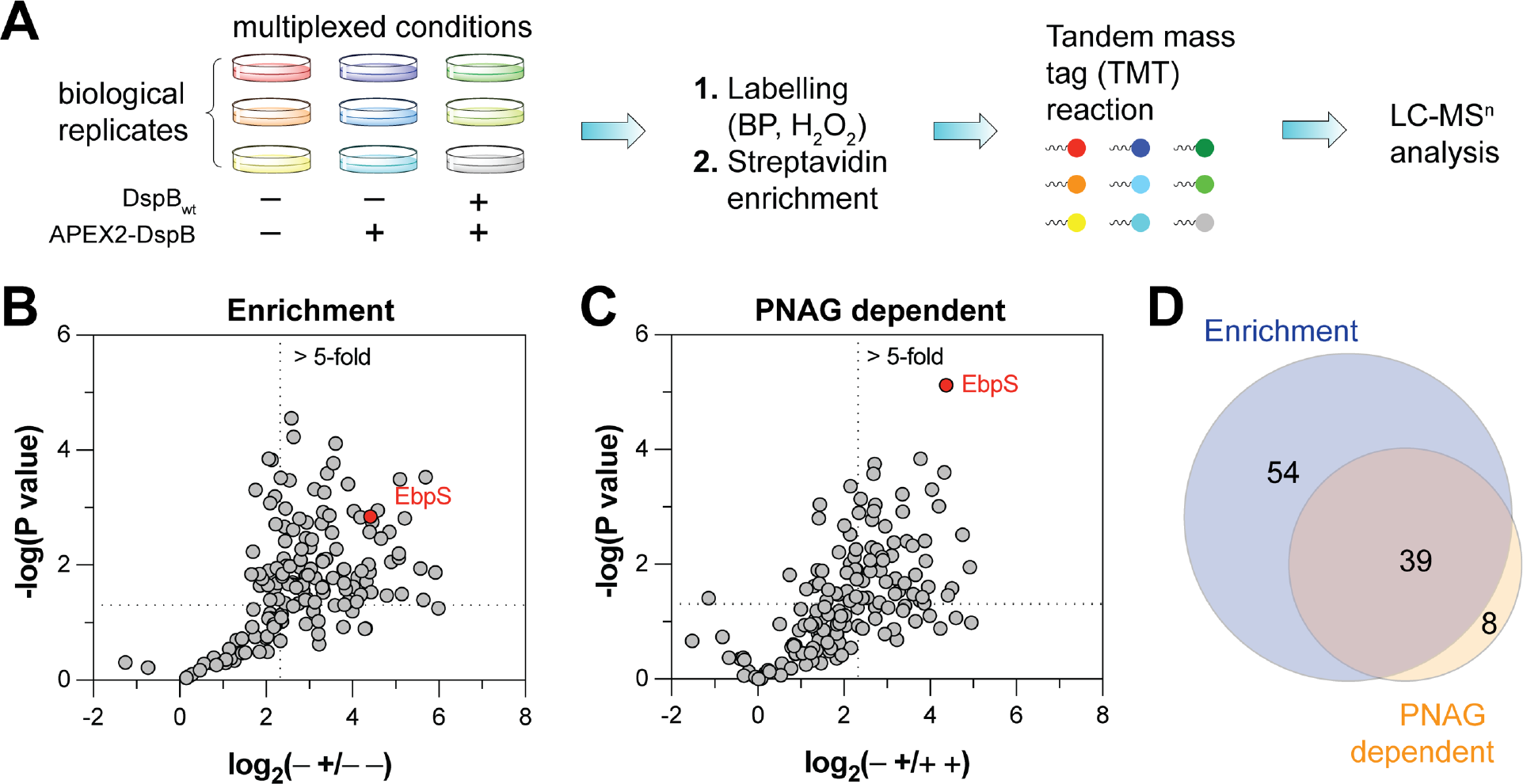
Quantitative proteomic analysis of biotinylated proteins isolated from *S. epidermidis* biofilm live cell proximity labeling experiments. (A) Experimental workflow for proximity labeling of the PNAG interactome of *S. epidermidis* biofilms and preparation of tandem mass tag-labeled tryptic peptides for analysis by LC-MS^n^. (B) Statistically significant (p < 0.05) and enriched (>5-fold) proteins found in biofilm samples treated with APEX2-DspB (– +) alone when compared to biofilm samples treated only with buffer (– –). (C) Statistically significant (p < 0.05) and enriched (>5-fold) proteins found in biofilm samples treated with APEX2-DspB (– +) alone when compared to biofilm samples treated with APEX2-DspB and DspB_wt_ to degrade PNAG. (D) A total of 93 proteins were found to be significantly enriched in biofilm samples incubated with APEX2-DspB compared to buffer alone, and 39 of these were significantly enriched in biofilms labeled with APEX2-DspB compared to those labeled with APEX2-DspB after digestion of PNAG by DspB_wt_ (i.e., enrichment is PNAG-dependent). Statistical significance was determined using a Welch’s multiple t-test as implemented in GraphPad Prism 10 (GraphPad Software, Boston, MA).

A total of 176 unique *S. epidermidis* proteins were identified across the three biological replicates (Fig. 2B–D, Table 1, and *SI appendix*). Of these, 93 were found to be significantly enriched (p < 0.05) by >5-fold in (– +) treated *S. epidermidis* biofilms compared to control (– –) biofilms (Fig. 2B). To identify proteins whose biotinylation is dependent on PNAG (Fig 2C), we also compared proteins isolated from (– +) biofilms to those from (+ +) in which PNAG had been hydrolyzed prior to labeling. A total of 47 proteins were significantly enriched by >5-fold in the (– +) relative to (+ +) samples, 39 of which were also enriched relative to the (– –) samples (Fig. 2D). These proteins represent putative PNAG-interacting proteins and are summarized in Table 1.

**Table 1.**
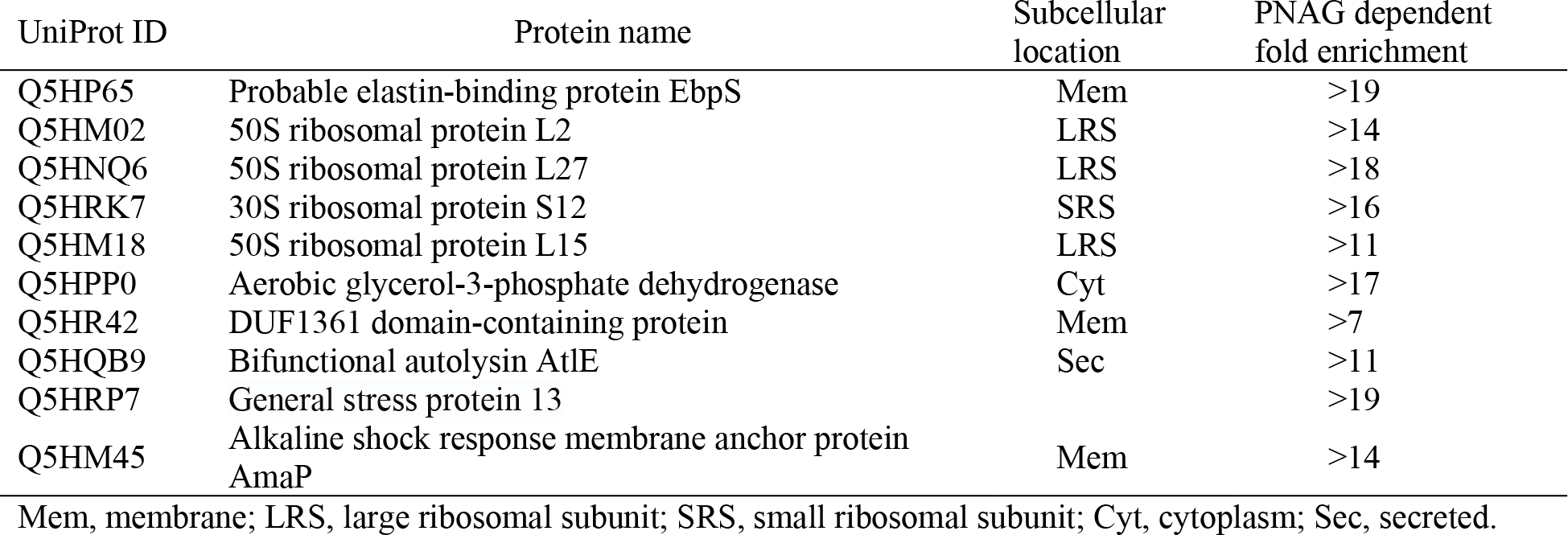
Select PNAG-interacting proteins identified by live cell proximity labeling.

The largest group of proteins identified consists of proteins associated with the 50S and 30S ribosomal subunit. It is not necessarily surprising to find ribosomal proteins enriched in a PNAG-dependent fashion as *S. aureus* has been shown to utilize environmental RNA within its biofilm EPS.^43^ The accumulation of environmental RNA in *S. aureus* biofilm is dependent on PNAG production and binds through ionic interactions with cationic PNAG.^43^ It seems likely that the ribosomal proteins identified in this study are released into the environment as a result of cell lysis during biofilm growth and maturation where they associate with PNAG through ionic interactions.

In addition to ribosomal proteins, known *S. epidermidis* biofilm proteins were also identified using our proximity labeling approach, including accumulation associated protein (Aap) and the bifunctional autolysin (AtlE). Aap is reported to contribute to biofilm formation through the formation of amyloid fibrils within the biofilm EPS.^44–47^ More recent reports also suggest the lectin-like A domain of Aap also contributes to initial cell adhesion during biofilm formation.^48–50^ While we observed a >8-fold enrichment of Aap through proximity labeling with APEX2-DspB, this enrichment was not dependent on PNAG. These results were consistent with Aap and PNAG playing independent roles during biofilm formation.^44,48,50–52^

AtlE is the major autolysin of *S. epidermidis* responsible for cell separation during cell division.^53–55^ The protein consists of both an amidase domain and a glucosaminidase domain along with 7 GW domains that function as cell wall-targeting modules.^56^ AtlE is primarily thought to contribute to biofilm formation through the release of extracellular DNA (eDNA),^57^ although a direct role for AtlE as an adhesin recognizing host extracellular matrix proteins has also been reported.^55^

The most significant protein enriched in a PNAG-dependent fashion from *S. epidermidis* biofilms was elastin-binding protein (EbpS) (Fig. 2C). The *S. aureus* homolog of EbpS is a member of the microbial surface components recognizing adhesive matrix molecules (MSCRAMM) family of proteins that contributes to cell adhesion through binding to human elastin peptides.^58,59^ However, subsequent studies indicate that *S. aureus* cells lacking EbpS retain the ability to bind elastin, suggesting there may be alternative roles for EbpS in cell adhesion and biofilm formation.^60,61^

EbpS in *S. epidermidis* is a 460 amino acid protein that is predicted to contain an intrinsically disordered *N*-terminal domain, followed by two hydrophobic patches, a single pass transmembrane (TM) helix, and a *C*-terminal lysin motif (LysM) domain (Fig. 3A). EbpS homologs are widely distributed amongst Staphylococcaceae. Interestingly, the *N*-terminal TNSHQD motif that was proposed to be essential for elastin recognition by *S. aureus* EbpS,^59^ is not conserved amongst EbpS homologs and is absent in the *S. epidermidis* EbpS homolog (Fig. S2). Conversely, the LysM domain is highly conserved in EbpS homologs.

**Figure 3.**
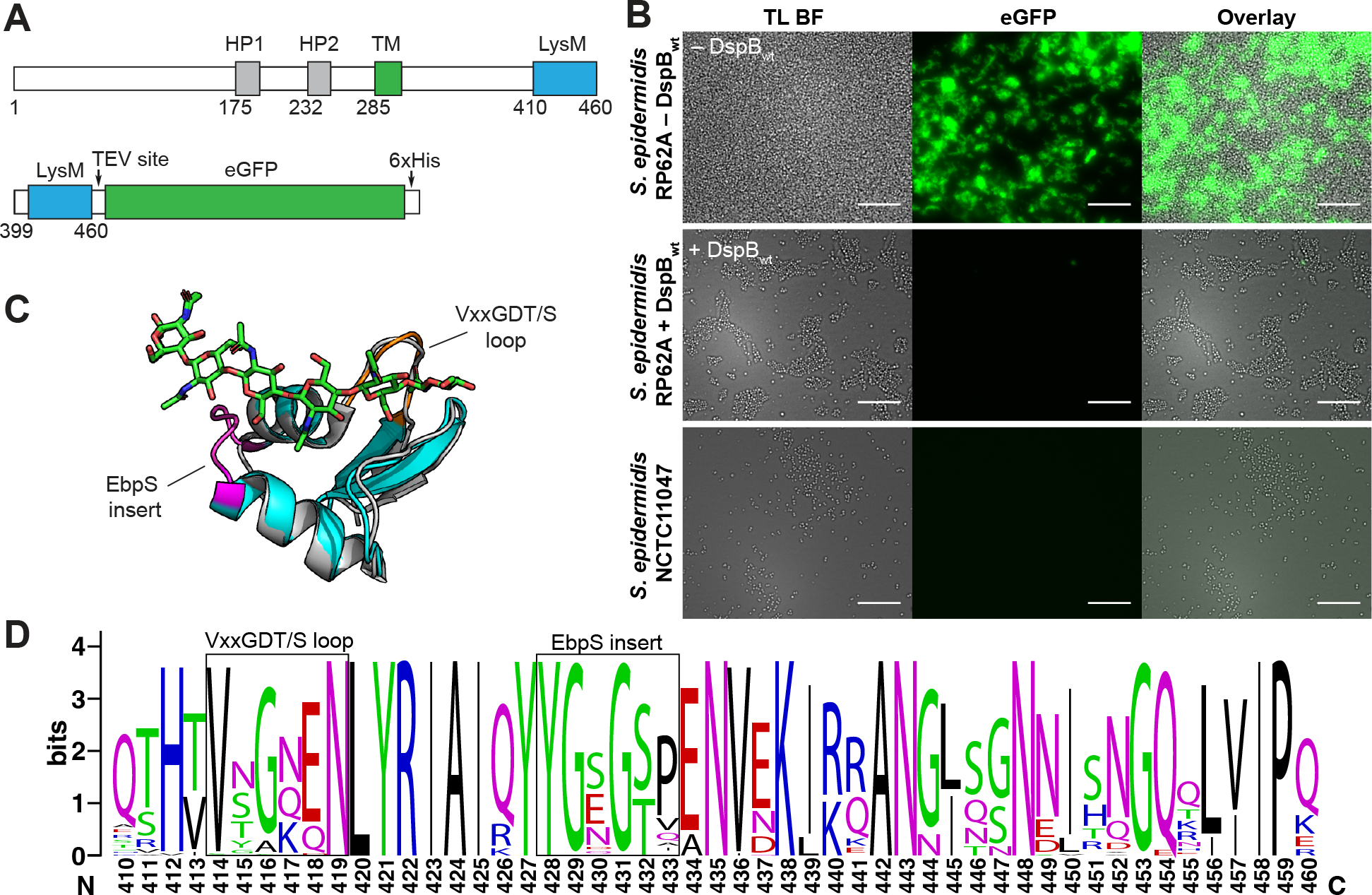
Evaluation of EbpS binding to PNAG in *S. epidermidis* biofilms. (A) The architecture of EbpS from S. *epidermidis* RP62A showing the conserved hydrophobic patches (HP1, HP2) transmembrane helix (TM) and *C*-terminal LysM domain (above), and a recombinant construct containing the EbpS LysM domain fused to eGFP used to evaluated binding to PNAG in *S. epidermidis* biofilms. (B) Fluorescent microscopy of *S. epidermidis* RP62A (PNAG positive) or NCTC11047 (PNAG negative, bottom) cells bound to EbpS LysM-eGFP construct before (top) or after (middle) hydrolysis of PNAG by DspB_wt_ (Scale bar = 20 µm). Binding of LysM-eGFP to *S. epidermidis* cells is PNAG dependent. (C) An overlay of an AlphaFold2 predicted structure of the EbpS LysM domain (cyan and magenta) compared to the LysM1 domain of the NlpC/P60 D,L-endopeptidase enzyme of *Thermus thermophilus* (PDB 4UZ3, gray) bound to a chitohexose oligosaccharide (green). The conserved EbpS insert (Y428–T432) is shown in magenta. (D) WebLogo showing conservation of the EbsS LysM domain amongst Staphylococcaceae. The position of a conserved EbpS insert, and non-conserved VxxGDT/S loop motif are indicated in boxes.

LysM domains are relatively small, typically 40 to 65 residue long, carbohydrate-binding motifs that are frequently found within enzymes responsible for hydrolysis of peptidoglycan or chitin,^62^ but have also been observed in plant receptor kinase proteins.^63^ LysM domains function as peptidoglycan-binding modules that specifically recognize *N*-acetylglycosamine (GlcNAc) residues present in peptidoglycan and chitooligosaccharides.^62^ The EbpS LysM domain is not thought to be surface exposed, but instead bound to cell wall peptidoglycan.^64^ However, given the similarity between PNAG (poly-(1→6)-β-GlcNAc) and chitin (poly-(1→4)-β-GlcNAc), we sought to evaluate if the LysM domain of *S. epidermidis* EbpS in fact functions as a PNAG-binding protein.

### Validation of PNAG binding by the EbpS LysM domain

We genetically fused the *C*-terminal LysM domain of EbpS (399–460) to eGFP through a short tobacco etch virus protease (TEV) cleavable linker (Fig. 3A). The resulting EbpS_LysM_-eGFP fusion protein was produced recombinantly in *E. coli*, and then used to measure binding to PNAG when added exogenously to *S. epidermidis* biofilms using fluorescent microscopy. EbpS_LysM_-eGFP effectively binds to *S. epidermidis* RP62A biofilms (Fig. 3B), but this binding can be disrupted by incubating the cells with DspB_wt_ to hydrolyze PNAG prior to binding to EbpS_LysM_-eGFP. Addition of EbpS_LysM_-eGFP to PNAG negative *S. epidermidis* NCTC11047, which does not form biofilms, showed no binding between EbpS_LysM_-eGFP and *S. epidermidis* NCTC11047 cells. Finally, when we incubated *S. epidermidis* RP62A biofilms with increasing concentrations of EbpS_LysM_-eGFP prior to treatment with DspB_wt_, we found that the EbpS_LysM_-eGFP protein is able to partially inhibit PNAG hydrolysis in a concentration dependent fashion (Fig. S3), suggesting that EbpS_LysM_-eGFP and DspB_wt_ bind to the same target. Taken together, these results demonstrate that the Ebps_LysM_ domain does not bind peptidoglycan and instead recognizes PNAG.

When we predict the structure of EbpS using AlphaFold 2 (AF2),^65,66^ only the TM domain and *C*-terminal LysM domain have predicted secondary structure (Fig. 3C and Fig. S4). The AF2 structure of EbpS_LysM_ aligns with the LysM domain of *Thermus thermophilus* NlpC/P60 D,L-endopeptidase enzyme bound to chitohexose^67^ with an RMSD of 0.542 Å. Similar results are seen when aligned to other known LysM domains bound to chitooligosaccharide ligands (Fig. S4). The major difference is that EbpS_LysM_ contains an additional five amino acid insertion between the two alpha helices (Fig. 3D) and lacks a highly conserved VxxGDT/S motif in the loop connecting the first beta sheet and alpha helix. This insertion is highly conserved in EbpS homologs but absent from the LysM domains of proteins co-crystalized with chitooligosacharide ligands. The conservation of this insert sequence in EbpS homologs and the lack of VxxGDT/S motif may play a role in altering the ligand specificity of EbpS to favor PNAG binding.

## Conclusions

PNAG is an important adhesive factor in biofilm formation of several Gram-positive and -negative human pathogens, yet little is known about the specific binding interactions between PNAG and other macromolecules within the biofilm EPS. Herein, we have developed a quantitative live cell proximity labeling method using a fusion protein of APEX2 with a catalytically inactive DspB_E184Q_ mutant to map the PNAG interactome in live *S. epidermidis* biofilms. Our results demonstrate that many cytosolic proteins, like ribosomal proteins that are anionic, closely associate with PNAG in live *S. epidermidis* biofilms. Presumably these proteins are released as a result of cell lysis occurring during biofilm maturation and remain associated with the cationic PNAG polysaccharides within the biofilm EPS. We also found that the known *S. epidermidis* biofilm protein Aap does not interact directly with PNAG. Importantly, we identified EbpS as a protein that was highly enriched in a PNAG-dependent fashion from *S. epidermidis* biofilms. We found that the LysM domain of EbpS does not recognize peptidoglycan but instead binds to PNAG. EbpS homologs are highly conserved amongst Staphylococcaceae, suggesting that EbpS binding to PNAG may be a general feature of Staphylococcaceae biofilms. Further work is required to characterize the structural basis for this interaction and elucidate its functional role in biofilm development. Finally, the live cell proximity labeling method reported here is not limited to the study of *S. epidermidis* biofilms but will be a powerful tool to study the PNAG interactome in the ever-increasing list of PNAG-producing bacterial species and provide critical insight into protein– carbohydrate binding interactions that are critical for biofilm formation.

## Materials and Methods

### General

All chemicals were purchased as analytical or reagent grade and used without further purification unless otherwise noted. Cells were grown in sterile autoclaved media. DspB_wt_ was prepared as previously described.^68^ APEX2-DspB, and EbpS_LysM_-eGFP were prepared as recombinant proteins in *E. coli* BL21(DE3) cells and purified using immobilized metal ion affinity chromatography (IMAC) as described below. Sequencing grade trypsin was obtained from Promega (Promega; V5111). All fluorescent microscopy measurements were performed on an Echo Revolve R3 fluorescent microscope (Echo, San Diego, CA) equipped with a 5-megapixel CMOS monochrome camera for fluorescent imaging, and a 12-megapixel color camera for brightfield imaging. All fluorescent imaging applications used LED illumination using either a FITC color cube (λ_ex_ = 470 nm, λ_em_ = 525 nm, cut off filter 495 nm) or TxRED color cube (λ_ex_ = 560 nm, λ_em_ = 630 nm, cut off filter 585 nm) and an Olympus 40× Fluorite Phase objective (numerical aperture 0.75, working distance 0.51 mm).

### APEX2-DspB protein preparation

A gene encoding APEX2 fused to the *N*-terminus of DspB_E184Q_ via a GSGGGGS linker, referred to as APEX2-DspB, was synthesized and codon optimized for expression in *E. coli* by GenScript. This gene was cloned into the NdeI and XhoI cut sites of a pET28a(+) vector to enable recombinant production of APEX2-DspB containing a *N*-terminal hexa-histidine tag. Recombinant APEX2-DspB was prepared by expressing the plasmid in *E. coli* BL21(DE3) cells. Cells from overnight cultures grown in Luria-Bertani (LB) broth containing 50 µg/mL kanamycin were used to inoculate a fresh 500 mL culture of terrific broth (TB) media containing 50 µg/mL kanamycin. Cells were grown to an OD_600_ = 0.6 at 37 °C in a shaking incubator, followed by induction with 0.5 mM isopropyl β-D-1-thiogalactopyranoside (IPTG). Cultures were then grown for 18 h at 18 °C before harvesting the cells by centrifuging for 20 min at 5000 × *g* and 4 °C.

The resulting cell pellet was resuspended in resuspension buffer (25 mM sodium phosphate, 150 mM sodium chloride, pH 7.4) containing 10 mM imidazole, and lysed by sonication. The cell lysate was clarified by centrifugation at 12,000 × *g* for 20 min at 4 °C, then purified by IMAC using a 1 mL HisTrap fast flow column and eluted in resuspension buffer containing 500 mM imidazole. Protein presence in the eluted sample was confirmed by SDS-PAGE analysis. The protein was exchanged into PBS buffer and concentrated to 100 mM using an Amicon 10,000 centrifugal filter device and stored at –80°C until use.

Heme was reconstituted into recombinant APEX2-DspB by incubating with hemin-HCl at a 2:1 molar ratio of hemin to protein for 30 min at room temperature. The excess hemin was removed by passing the protein sample over a column of DEAE-cellulose and then purifying the resulting protein by size exclusion chromatography on a Sephacryl S-200 column eluting with 30 mL of resuspension buffer. The presence of heme in the protein was confirmed by examining the ratio of absorbance at 405 nm to total protein absorbance at 280 nm. A ratio ≥ 1.5 (heme:protein) was generally considered to be sufficient for use in live cell proximity labeling.

### Growth of *S. epidermidis* biofilms

*S. epidermidis* strain RP62A (ATCC 35984) or NCTC11047 (ATCC 14990) cells were grown in an overnight culture (18 hrs) in tryptic soy broth (TSB) media. The cells were then diluted 1:10 in freshly prepared TSB media and gently shaken for 5 min at 37 C. Biofilms were then grown in 6-well microtiter plates by transferring 7 mL of culture to each well, covering the plate, and incubating at 37 °C in static culture for 48 h.

### Proximity labeling of live *S. epidermidis* biofilms

Biofilms after 48 h growth were subjected to the following labeling process in duplicate. Replicate biofilms were incubated in 300 µL phosphate buffer (100 mM NaCl and 25 mM Na_2_HPO_4_, pH 6.0) containing 2 µM of DspB-APEX2 enzyme (– + samples) and incubated for 30 min at 37 °C with gentle shaking. Separate biofilms samples were incubated with the same volume of phosphate buffer containing both 2 µM DspB-APEX2 and 2 µM DspB_wt_ enzyme (+ + samples). Finally, additional replicate biofilms were incubated for the same period in phosphate buffer alone (– – samples). Following the 30-min incubation, all biofilm samples from all three sets of conditions were washed with 300 µL of phosphate buffer for three times and centrifuged to pellet the cells after each wash step. The resulting samples were then incubated with 2.5 mM BP in 300 µL of phosphate buffer for 30 min at 37 °C. The excess BP mixture was removed before incubating the biofilms in new 300 µL of phosphate buffer containing 0.5 mM H_2_O_2_. The biotinylation was allowed to proceed for 1 min before being quenched by adding an equal volume of quench solution containing (5 mM Trolox, 10 mM sodium ascorbate, and 10 mM sodium azide in phosphate buffer). All samples were then washed with quench solution three additional times. Biofilm cells were flash frozen and stored at –80 °C prior to analysis. The entire labeling procedure was repeated to generate a minimum of three biological replicates for each sample condition.

### Fluorescent labeling of biotinylated biofilms with Streptavidin-TexasRed

Biotinylated biofilm samples used for fluorescence imaging were incubated with a 1:2000 dilution of streptavidin-TexasRed conjugate (300 µL volume) in the dark at room temperature for 30 min to label biotinylated proteins. The biofilm samples were washed with buffer A before imaging. Labeled biofilm cells were immobilized on an agarose pad prepared as described previously,^69^ by pipetting the cells between the agarose pad and glass cover slip immediately prior to imaging.

### Preparation and enrichment of biotinylated proteins

Biotinylated biofilm samples were re-suspended in 1 × RIPA buffer (50 mM Tris-HCl, pH 8 with 150 mM NaCl, 5 mM EDTA, 0.1% SDS, 1% Triton X-100, 5 mM Trolox, 10 mM sodium ascorbate, 10 mM sodium azide) containing 1 × protease inhibitor cocktail (Sigma), 1 µM lysostaphin and 1 µM hen egg white lysozyme. The cells were incubated for 1 h at 37 °C, and then frozen at –80 °C overnight. The cells were subsequently subjected to 5 × freeze thaw cycles by incubating cells in a water bath at 42 °C for 5 min, followed by freezing in a dry ice bath for 5 min. The cell debris was removed by centrifugation at 17,000 × *g* for 45 min at 4 °C, and the supernatant containing extracted proteins was collected. The protein concentration in each sample was determined using the bicinchoninic acid (BCA) assay and the volumes were adjusted to contain 1 mg/mL total protein.

For Western blot analysis, 50 µL of each protein samples were incubated with 50 µL of DynaBeads MyOne Streptavidin C1 beads (Thermo Fisher Scientific; 65001) that had been pre-incubated in PBS buffer containing 5% bovine serum albumin (BSA) for 30 min at room temperature. The beads were subsequently washed with 500 µL of PBS buffer containing 0.1% SDS and 1% BSA × 3. Proteins were eluted by heating the beads in 100 µL of 1 × Laemmli-buffer to 95 °C for 5 min. Equal volumes of input and eluted protein samples were then analyzed by SDS-PAGE, imaged using stain free imaging (BioRad), and then subjected to Western blot analysis using a streptavidin-HRP conjugate for detection. The blot was incubated with streptavidin biotin conjugate for 1 hr followed by washing with PBST three times before detection. The blot was imaged via chemiluminescent detection using Clarity Western ECL substrate (BioRad) according to the manufacturer’s directions and imaged using an iBright CL1000 imaging system (Thermo Scientific).

For quantitative proteomics analysis, the protein samples were further aliquoted (100 µL containing 300 µg of proteins in 100 mM triethylammonium bicarbonate (TEAB) buffer for enrichment, protein digestion, and TMT^10^-plex TMT labeling (Thermo Fisher Scientific; catalog no. 90113) according to directions of the manufacturer. Briefly, first protein disulfide bonds were reduced using 5 µL of 200 mM tris(2-carboxyethyl)phosphine (TCEP) and incubated for 1 h at 55 °C, then alkylated with 5 µL of 375 mM iodoacetamide and incubated at room temperature protected from light for 30 min. Next, the reduced proteins were enriched by incubating with DynaBeads MyOne Streptavidin C1 beads (Thermo Fisher Scientific; catalog no. 65001). Beads were washed three times with filtered 1 × PBS buffer before enrichment. The resulting proteome was incubated with the beads for 30 min at room temperature. Unbound proteins (supernatant) were separated using a magnet, and the beads with biotinylated proteins were further washed 3 times with PBS buffer. The samples were then subjected to on-bead digestion by incubating with trypsin (1:40 ratio by mass) at 37 °C overnight (⁓18 h). The samples were centrifuged, and the supernatants were transferred to sterile Low Bind microcentrifuge tubes (TermoFisher).

Peptide concentrations were determined using a Take3 micro-volume plate (Agilent Technologies, Santa Clara, CA) before labeling each sample with the appropriate TMT^10^-plex reagent. All samples were diluted to the same protein amount (5.6 µg). TMT reagents were added in a ratio of 41 µL for every 100 µg of digested protein and samples were incubated for 1 h at room temperature, then quenched with 8 µL of 5% hydroxylamine and incubated for 15 min at room temperature. A total of 9 samples were prepared in this way, three biological replicates for each of the three biotinylation conditions. These barcoded samples were combined into one new microcentrifuge tube and vacuum-dried in a speedvac. The combined sample was desalted using a Pierce peptide desalting spin column (Thermo Fisher Scientific; catalog no. 89852) and vacuum-dried in a Centrivap (Labconco, Kansas City, MO).

### Quantitative proteomics by LC-MS^n^

The sample was analyzed in technical triplicate by quantitative LC-MS^z^. The TMT^10^-plex-tagged sample (1 µg) was loaded onto an Acclaim PepMap100 C18 trap column (Thermo Fisher Scientific, 100 µm × 2 cm, 100 Å, 5 µm) at 5 µL/min using an UltiMate 3000 RSLCnano system (Thermo Fisher Scientific) prior to separation on a micropillar array column (µPAC; Thermo Fisher Scientific, 200 cm, 50 °C) at 600 nL/min using a 240-min gradient consisting of Buffer A (0.1% formic acid in MS-grade water) and Buffer B (0.1% formic acid in MS-grade acetonitrile) as follows: starting at 1% B for 5 min; increased to 7% B over 15 min; to 25% B over 115 min; to 32% B over 25 min; to 45% B over 33 min; to 75% B over 7 min; then eluent composition was held at 75% B for 8 min; before reducing to 2% B over 2 min; column equilibration took place at 2% B for 30 min.

Peptides were analyzed with electrospray ionization (ESI) and detected on quadrupole-orbitrap-ion trap tribrid mass spectrometer (Orbitrap Fusion Lumos, Thermo Fisher Scientific). Survey full (MS^1^) scans were acquired in the positive ion mode in the Orbitrap analyzer between *m/z* 400–1600 with a spectral resolution of 120,000 full width at half maximum (FWHM) using a standard automatic gain control (AGC) target and the auto maximum injection time mode. For protein identification, precursor ions with charges +2–+6 and intensities > 5.0 × 10^3^ counts were selected for MS^2^ using data-dependent acquisition with a 2.5 s cycle time and 60 s dynamic exclusion. MS^2^ employed collision-induced dissociation (CID) in the ion trap with 35% normalized collision energy (NCE) using turbo ion trap scan rate, auto maximum injection time and scan range modes, standard AGC target and isolation window of 0.7 m/z. For quantification, 10 of the MS^2^ fragment ions were isolated by synchronous precursor selection (SPS) for higher-energy collisional dissociation (HCD) at 55% NCE. The resulting fragment ions (MS^3^ spectra) were analyzed in the Orbitrap between *m/z* 100–500 with a resolution of 50,000 FWHM and a 0.7 m/z isolation window (standard AGC target, auto maximum injection time mode). Precursors with TMT tag loss were excluded from MS^3^ selection to maximize quantification from TMT-labeled peptides.

### Proteomics Data Analysis

The primary MS files were analyzed in Proteome Discoverer version 2.2 (Thermo Fisher Scientific). Technical replicates were combined and searched against the UniProt *S. epidermidis* strain RP62A reviewed and unreviewed proteome (2,492 sequences, downloaded on 4/25/2023) and a database of common contaminants using SEQUEST-HT. The following dynamic modifications were included in the search: TMT^10^-plex (229.163 Da) on peptide N-termini and lysine residues, BP labeling of tyrosine residues (361.460 Da), biotinylation of lysine residues (226.078 Da), oxidation of methionine residues, and acetylation of the protein *N*-termini. Carbamidomethylation on cysteine residues was included as a fixed modification. Maximum two trypsin missed cleavages were allowed in the search. Mass tolerances were <10 ppm for precursor ions and 0.6 Da for fragment ions. Peptide and protein identifications were both filtered to < 1% false discovery rate (FDR).

A TMT^10^-plex quantification method was customized for the 9 TMT channels used in this experiment and the corresponding TMT lot-specific isotopic impurity corrections. Reporter ions were quantified on the bases of ion signal intensity with a signal-to-noise ratio of ≥ 30. The reporter ion intensities were normalized to the total peptide amount in Proteome Discoverer. Only Master Proteins with < 1% FDR were reported as identified, excluding contaminants. The MS proteomics data have been deposited to the ProteomeXchange Consortium via the PRIDE^70^ partner repository with the dataset identifier PXD044531.

For subsequent data analysis, the relative ion abundances for individual identified proteins were normalized to the ion abundance for pyruvate carboxylase (Uniprot Q5HQ53), a natively biotinylated protein for each of the nine samples. The statistical significance of protein enrichment was determined by comparing the log_2_-transformed normalized protein abundance from three biological replicates using a Welch’s multiple t-test as implemented in GraphPad Prism 10 (GraphPad Software, Boston, MA). Proteins that were significantly (p < 0.05) enriched by >5-fold were considered as hits. Enrichment was measured for (– +) samples relative to both control (– –) and DspB_wt_ treated samples (+ +).

### Cloning of EbpS C-terminal LysM domain as an eGFP fussion

A gene encoding *S. epidermidis* EbpS was synthesized, and codon optimized for expression in *E. coli* by Genscript. The sequence encoding the *C*-terminal LysM domain (401-460) was subsequently subcloned into a pET28a(+) vector containing a *C*-terminal eGFP and 6 × His tag connected via a short TEV protease cleavable linker (pLV28). The sequence was verified by single pass Sanger sequencing and used to transform *E. coli* BL21(DE3) cells.

*E*.*coli* BL21 cells containing the pLV28 plasmid were grown in overnight cultures in LB broth containing 50 µg/mL kanamycin. 10 mL of overnight culture was used to inoculate a fresh 500 mL LB broth containing 50 µg/mL kanamycin, which was grown at 37 °C with shaking until reaching an OD_600_ = 0.6. The culture was induced via the addition of 0.5 mM IPTG and grown for 18 h at 18 °C. The cells were harvested by centrifugation at 5000 × *g* for 20 min at 4 °C. The harvested pellet was resuspended in resuspension buffer containing 10 mM imidazole and lysed via sonication. The cell lysate was clarified by centrifugation at 12,000 × *g* for 20 min at 4 °C, then purified by IMAC using a 1 mL HisTrap fast flow column and eluted in resuspension buffer containing 500 mM imidazole. The protein was exchanged into PBS buffer, concentrated using an Amicon 10,000 MWCO centrifugal filter device, and stored at –80 °C until use.

### Imaging LysM-eGFP binding *S. epidermidis* cells

*S. epidermidis* RP62A and NCTC11047 biofilms were grown as described above for 48 h. Biofilm cells were then incubated with either phosphate buffer alone or phosphate buffer containing 2 µM DspB_wt_ for 1 hr at 37 °C. Both DspB_wt_ treated and untreated biofilm samples were then incubated with varying concentrations of EbpS_LysM_-eGFP fusion protein between 1–15 µM. The samples were then washed with phosphate buffer 3 × to remove any unbound EbpS_LysM_-eGFP and immobilized on an agarose pad for analysis by fluorescent microscopy. Cells were resuspended in 100 µL of PBS buffer after washing, then 5 µL was transferred between the agarose pad and glass coverslip.

For measurement of EbpS_LysM_-eGFP inhibition of PNAG degradation by DspB_wt_, 48 hr old *S. epidermidis* biofilm samples were simultaneously incubated with DspB_wt_ (2 µM) and EbpS_LysM_-eGFP (1–15 µM) in phosphate buffer. Residual biofilm were then imaged as described above.

## Abbreviations

EPS: extracellular polymeric substance
VPS: vibrio polysaccharide
PNAG: poly-(1→6)-β-*N*-acetylglucosamine
LPS: lipopolysaccharide
WTA: wall teichoic acids
DspB: Dispersin B
APEX2: engineered ascorbate peroxidase
eGFP: enhanced green fluorescent protein
MS: mass spectrometry
HRP: horseradish peroxidase
TMT: tandem mass tag
LC-MS: liquid chromatography-mass spectrometry
MS^n^: tandem mass spectrometry
Aap: accumulation associated protein
AtlE: bifunctional autolysin
EbpS: elastin binding protein
LysM: lysin motif
eDNA: extracellular DNA

## Acknowledgements

This work was supported by the National Institute of General Medical Science of the National Institutes of Health under award number R35GM147110. The content is solely the responsibility of the authors and does not necessarily represent the official views of the National Institutes of Health.

