## Supplemental Information for "Mapping protein–exopolysaccharide binding interaction in *Staphylococcus epidermidis* biofilms by live cell proximity labeling"

### Supporting Information

1. Supplemental Table S1 2
2. Supplemental Figures S1–S4 3
3. Supplemental References 6

### Table S1. PNAG interacting proteins identified by live cell proximity labeling

| UniProt ID | Protein name | Subcellular location | PNAG dependent fold enrichment |
|---|---|---|---|
| Q5HP65 | Probable elastin-binding protein EbpS | Mem | >19 |
| Q5HM02 | 50S ribosomal protein L2 | LRS | >14 |
| Q5HM32 | 30S ribosomal protein S9 | SRS | >7 |
| Q5HNQ6 | 50S ribosomal protein L27 | LRS | >18 |
| Q5HM22 | 50S ribosomal protein L36 | LRS | >7 |
| Q5HRK7 | 30S ribosomal protein S12 | SRS | >16 |
| Q5HM18 | 50S ribosomal protein L15 | LRS | >11 |
| Q5HKT5 | Diacetyl reductase [(S)-acetoin forming] | Cyt | >5 |
| Q5HM15 | 50S ribosomal protein L18 | LRS | >7 |
| Q5HPP0 | Aerobic glycerol-3-phosphate dehydrogenase | Cyt | >17 |
| Q5HR42 | DUF1361 domain-containing protein | PM | >7 |
| Q5HQB9 | Bifunctional autolysin AtlE | Sec | >11 |
| Q5HM00 | 50S ribosomal protein L4 | LRS | >5 |
| Q5HPT4 | Elongation factor Ts | Cyt | >7 |
| Q5HQV1 | 2,3-bisphosphoglycerate-independent phosphoglycerate mutase | Cyt | >8 |
| Q5HR56 | Uncharacterized protein | | >22 |
| Q5HPV3 | 30S ribosomal protein S16 | SRS | >15 |
| Q5HNM4 | 50S ribosomal protein L35 | LRS | >11 |
| Q5HNX1 | 30S ribosomal protein S21 | SRS | >10 |
| Q5HRL4 | 50S ribosomal protein L1 | LRS | >12 |
| Q5HPS2 | Translation initiation factor IF-2 | Cyt | >6 |
| Q5HNM8 | Trigger factor | Cyt; LRS | >7 |
| Q5HNI5 | 30S ribosomal protein S4 | SRS | >9 |
| Q5HM97 | Fructose-bisphosphate aldolase | Cyt | >8 |
| Q5HPV8 | Ribonuclease 3 | Cyt | >8 |
| Q5HR71 | Probable transcriptional regulatory protein SERP0322 | Cyt | >6 |
| Q5HLZ9 | 50S ribosomal protein L3 | LRS | >15 |
| Q5HM31 | 50S ribosomal protein L13 | LRS | >24 |
| Q5HM04 | 50S ribosomal protein L22 | LRS | >8 |
| Q5HQ20 | Phenol soluble modulin beta 1 | Sec | >6 |
| Q5HM14 | 50S ribosomal protein L6 | LRS | >13 |
| Q5HLY6 | Lipid II:glycine glycyltransferase | Mem | >10 |
| Q5HQ75 | Pyruvate dehydrogenase E1 component subunit beta | | >5 |
| Q5HN52 | YtxH domain-containing protein | | >8 |
| Q5HM03 | 30S ribosomal protein S19 | SRS | >11 |
| Q5HM01 | 50S ribosomal protein L23 | LRS | >15 |
| Q5HQX7 | Ribosome hibernation promotion factor | Cyt; LRS; SRS | >20 |
| Q5HPG5 | Aminoacyltransferase FemB | Mem | >10 |
| Q5HMT6 | Uncharacterized protein | | >10 |
| Q5HM10 | 50S ribosomal protein L24 | LRS | >13 |
| Q5HRP7 | General stress protein 13 | | >19 |
| Q5HM45 | Alkaline shock response membrane anchor protein AmaP | Mem | >14 |
| Q5HKU2 | Arginine deiminase | Cyt | >6 |
| Q5HPB3 | Peptide methionine sulfoxide reductase MsrA | | >5 |
| Q5HNH1 | Smooth muscle caldesmon | Mem | >14 |
| Q5HRK6 | 30S ribosomal protein S7 | SRS | >7 |
| Q5HKM3 | Immunodominant antigen B, putative | | >14 |

Mem, membrane; LRS, large ribosomal subunit; SRS, small ribosomal subunit; Cyt, cytoplasm; Sec, secreted.

### Supplemental Figures

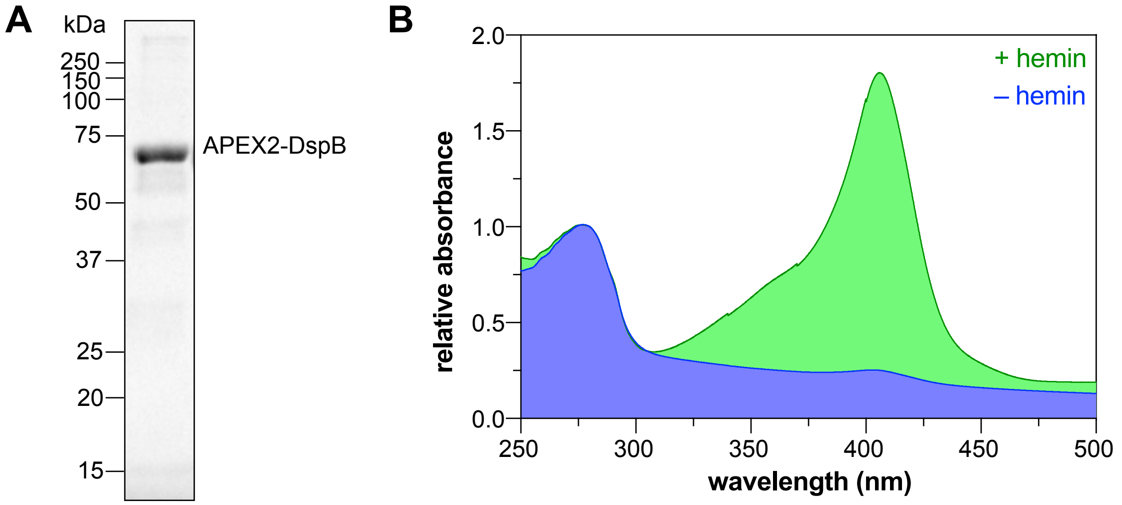


**Figure S1.** Preparation and analysis of recombinant APEX2-DspB protein. (A) SDS-PAGE showing purified APEX2-DspB protein used in these studies after heme reconstitution and purification by size exclusion chromatography. A major band at ~68 kDa corresponding to the APEX2-DspB fusion protein is observed. (B) UV-vis absorbance spectra for APEX2-DspB before (- hemin) and after (+ hemin) heme reconstitution. Spectra were recorded for the purified protein after size exclusion chromatography and are normalized relative to the 280 nm absorbance. A 405 nm/280 nm absorbance ratio >1.5 was deemed to be suitable for live cell proximity labeling applications.

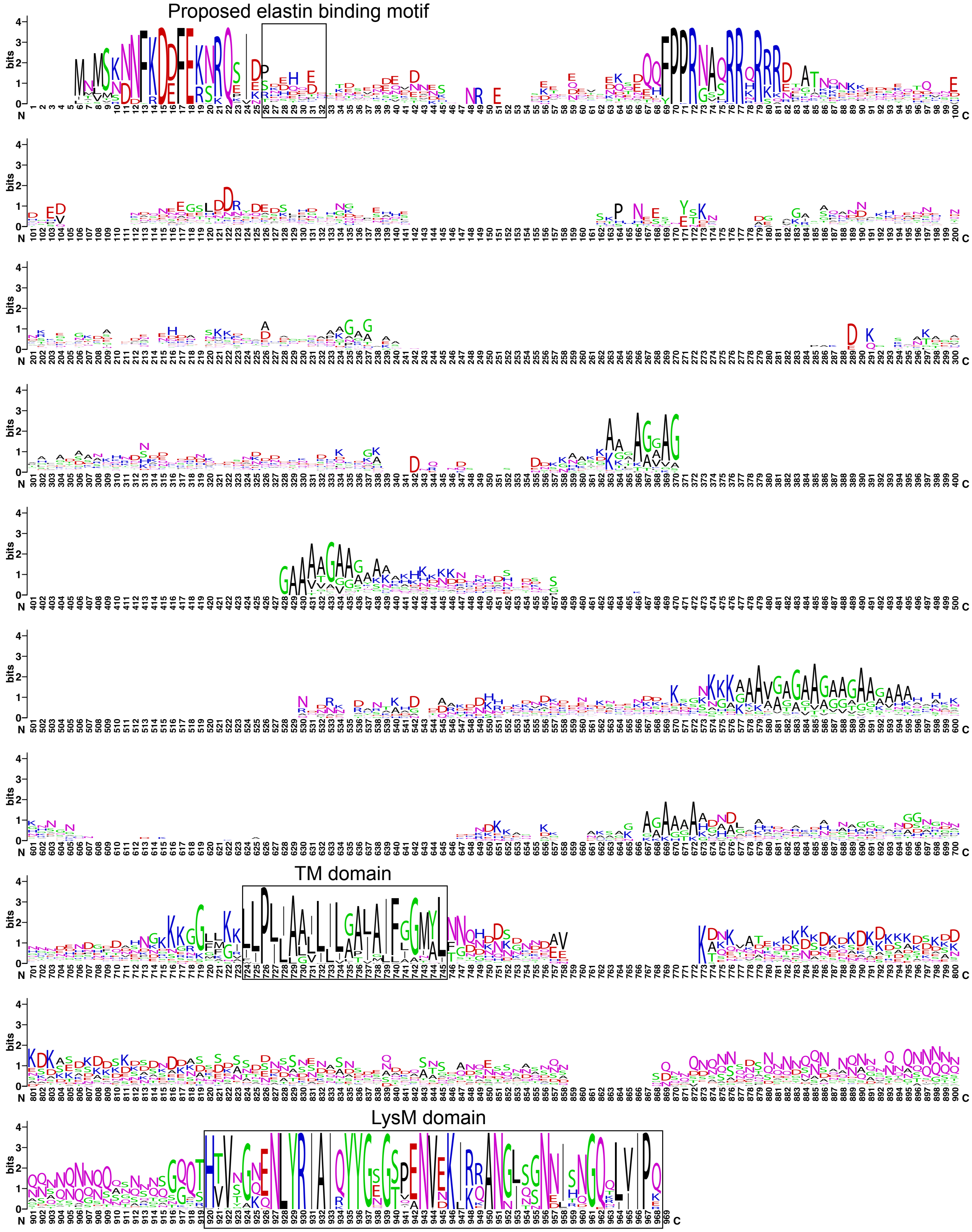


**Figure S2.** WebLogo diagram showing the sequence conservation of EbpS homologs from 51 species of Staphylococcaceae. The proposed elastin binding motif, transmembrane (TM) domain and LysM domain are indicated using boxes.

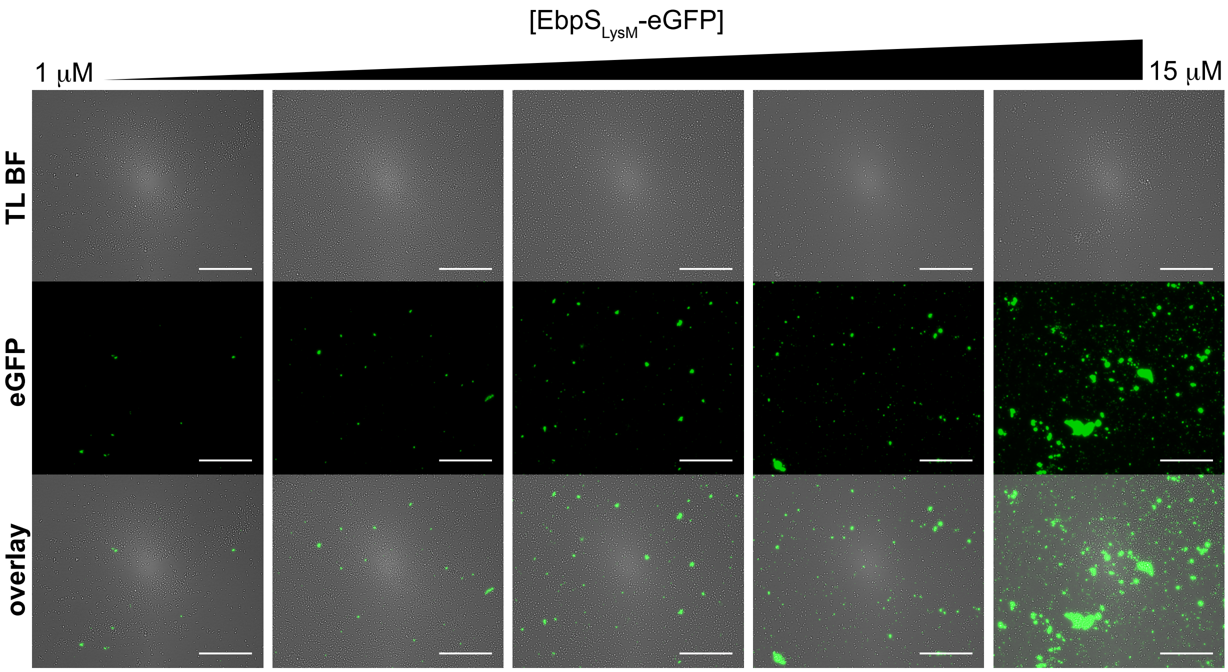


**Figure S3.** Competitive binding of EbpS$_{LysM}$-eGFP and DspB$_{wt}$ during *S. epidermidis* RP62A biofilm dispersal. Biofilms incubated with DspB$_{wt}$ (2 μM) and increasing concentrations of EbpS$_{LysM}$-eGFP (1 μM, 2 μM, 5 μM, 10 μM, and 15 μM) show more residual biofilm biomass remaining and greater PNAG remaining, indicating that EbpS$_{LysM}$-eGFP and DspB$_{wt}$ bind competitively to the same target (scale bar = 50 μm).

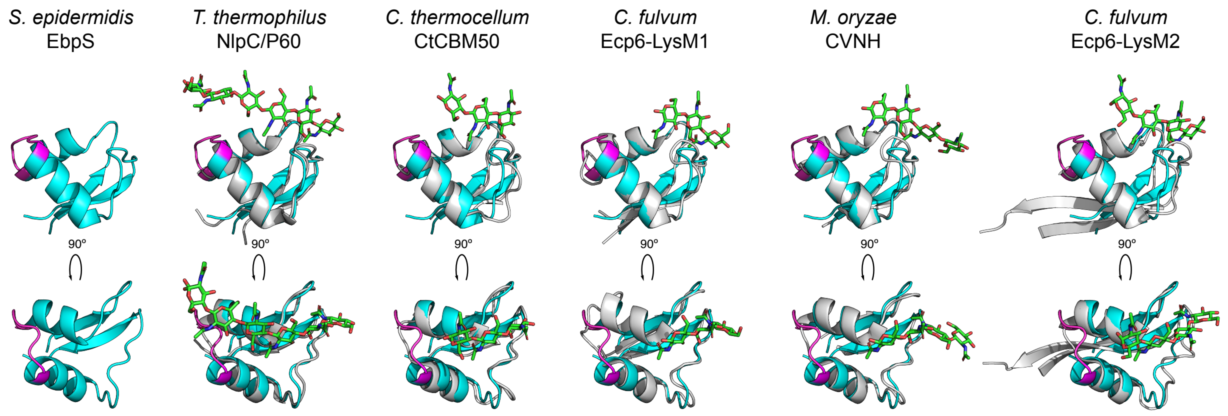


**Figure S4.** Comparison of the alphafold2 predicted structure of *S. epidermidis* EbpS$_{LysM}$ domain (cyan) to known LysM domain crystal structures (grey) that were co-crystalized with chitooligosaccharides (green), including *T. thermophilus* NlpC/P60 (PDB 4UZ3),[1] *C. thermocellum* CtCBM50 (PDB 7R1L), *C. fulvum* Ecp6-LysM1 (PDB 6Q40),[2] *M. oryzae* CVNH (PDB 5C8Q),[3] and *C. fulvum* Ecp6-LysM2 (PDB 4B9H).[4]
